# Mitigating autocorrelation during spatially resolved transcriptomics data analysis

**DOI:** 10.1101/2023.06.30.547258

**Authors:** Kamal Maher, Morgan Wu, Yiming Zhou, Jiahao Huang, Qiangge Zhang, Xiao Wang

**Affiliations:** Broad Institute of MIT and Harvard, Cambridge, MA, USA; Computational and Systems Biology Program, Massachusetts Institute of Technology, Cambridge, MA, USA; Stanley Center for Psychiatric Research, Broad Institute of MIT and Harvard, Cambridge, MA, USA; Department of Chemistry, Massachusetts Institute of Technology, Cambridge, MA, USA; McGovern Institute for Brain Research, Department of Brain and Cognitive Sciences, Massachusetts Institute of Technology, Cambridge, MA, USA

## Abstract

Several computational methods have recently been developed for characterizing molecular tissue regions in spatially resolved transcriptomics (SRT) data. However, each method fundamentally relies on spatially smoothing transcriptomic features across neighboring cells. Here, we demonstrate that smoothing increases autocorrelation between neighboring cells, causing latent space to encode physical adjacency rather than spatial transcriptomic patterns. We find that randomly sub-sampling neighbors before smoothing mitigates autocorrelation, improving the performance of existing methods and further enabling a simpler, more efficient approach that we call <u>sp</u>atial <u>in</u>tegration (SPIN). SPIN leverages the conventional single-cell toolkit, yielding spatial analogies to each tool: clustering identifies molecular tissue regions; differentially expressed gene analysis calculates region marker genes; trajectory inference reveals continuous, molecularly defined anatomical axes; and integration allows joint analysis across multiple SRT datasets, regardless of tissue morphology, spatial resolution, or experimental technology. We apply SPIN to SRT datasets from mouse and marmoset brains to calculate shared and species-specific region marker genes as well as a molecularly defined neocortical depth axis along which several genes and cell types differ across species.

## Introduction

The introduction of spatially resolved transcriptomics (SRT) has raised the fundamental question of how to leverage paired spatial and gene expression data^1–3^. A prominent approach is to combine this information to identify molecularly defined tissue regions, yielding insight into the transcriptional basis of tissue architecture and function^4–8^. While current methods for molecular region identification vary in their exact approaches, they each fundamentally rely on smoothing gene expression features over the tissue, i.e. setting each cell’s feature vector to a weighted sum of itself and its spatial nearest neighbors. For instance, UTAG^9^ implements smoothing through message passing, in which the weights used for neighborhood averaging correspond to the physical distances between the given cell and each neighbor. Similarly, STAGATE^10^ uses a graph convolutional network to smooth features over spatial nearest neighbors with learnable weights for each neighbor. Smoothing is fundamental to region identification tasks because it is equivalent to isolating low-frequency, large-scale patterns over the spatial neighbors graph^11^, which defines molecular regions. Alternatively, smoothing can be understood as a sliding window average over the tissue, blurring away variation in small-scale cell type-specific expression signals and instead revealing large-scale regional signals.

Here, we describe a fundamental limitation of smoothing that prevents the use of multiple existing region identification methods. Because spatially adjacent cells have almost identical neighborhoods, their smoothed expression features are also almost identical. In other words, smoothing increases autocorrelation between neighboring cells’ expression features. This leads to spatial reconstruction of the tissue in latent space: adjacency in latent space reflects adjacency in physical space. We show that autocorrelation can be mitigated via several strategies, including randomly sub-sampling each cell’s spatial neighborhood before smoothing. This allows adjacent cells to vary in their *exact* neighborhood compositions while maintaining their *general* molecular compositions, ultimately enabling the resulting latent space to represent meaningful spatial transcriptomic features rather than just the spatial adjacency of cells.

We further utilize subsampling to develop a simpler and more efficient alternative to existing molecular region characterization methods which we call <u>sp</u>atial <u>in</u>tegration (SPIN). SPIN leverages the conventional single-cell analysis toolkit to achieve spatial analogies to each tool: clustering identifies molecular tissue regions; differentially expressed gene (DEG) analysis calculates marker genes for each region; trajectory inference identifies continuous molecular trajectories in physical space; and integration allows each of these tools to be applied jointly across multiple datasets, regardless of tissue morphology, spatial resolution, or experimental technology. As this approach is based on the conventional single-cell toolkit, it does not require GPU acceleration and is as efficient as conventional single-cell analysis, with the exception of a fast preliminary smoothing step. We demonstrate this approach by comparing spatial transcriptomic features across several SRT brain datasets from mouse and marmoset species as well as across multiple experimental technologies, showing that datasets with arbitrary morphologies and spatial resolutions can be jointly spatially analyzed using conventional single-cell tools. We identify genes and cell types with spatial distributions that differ across species, most notably the widely expressed neuronal marker *Cck* as well as oligodendrocyte and interneuron cell types. We provide an implementation of SPIN based on Scanpy^12^ along with basic usage principles and a tutorial notebook here:

https://github.com/wanglab-broad/spin

## Results

### Smoothing reconstructs spatial connectivity in latent space

Single-cell transcriptomics analysis typically begins with molecular cell type clustering, during which single-cell gene expression vectors are stacked into a cell-by-gene matrix, followed by dimension reduction using principal components analysis (PCA), clustering using the Leiden^13^ algorithm, and visualization using uniform manifold approximation and projection (UMAP)^14^ (Fig. 1a). Current methods for molecular tissue region clustering follow a similar workflow, but with the addition of a preliminary smoothing step in which each cell’s expression feature is set to a weighted average of itself and its spatial nearest neighbors. To better understand these methods, we investigated a simplified model in which smoothing constitutes neighborhood averaging with equal weights across neighbors (Fig. 1b). We identified a potential issue with this approach in which physically adjacent cells have nearly identical neighborhoods, yielding smoothed expression features that are nearly equal (Fig. 1c). This can also be understood as smoothing-induced spatial autocorrelation. Because downstream methods such as Leiden and UMAP operate based on a nearest neighbors graph (in feature space), we reasoned that the resulting latent graph would represent the spatial proximity of cells to one another rather than the desired spatial transcriptomic features.

**Figure 1.**
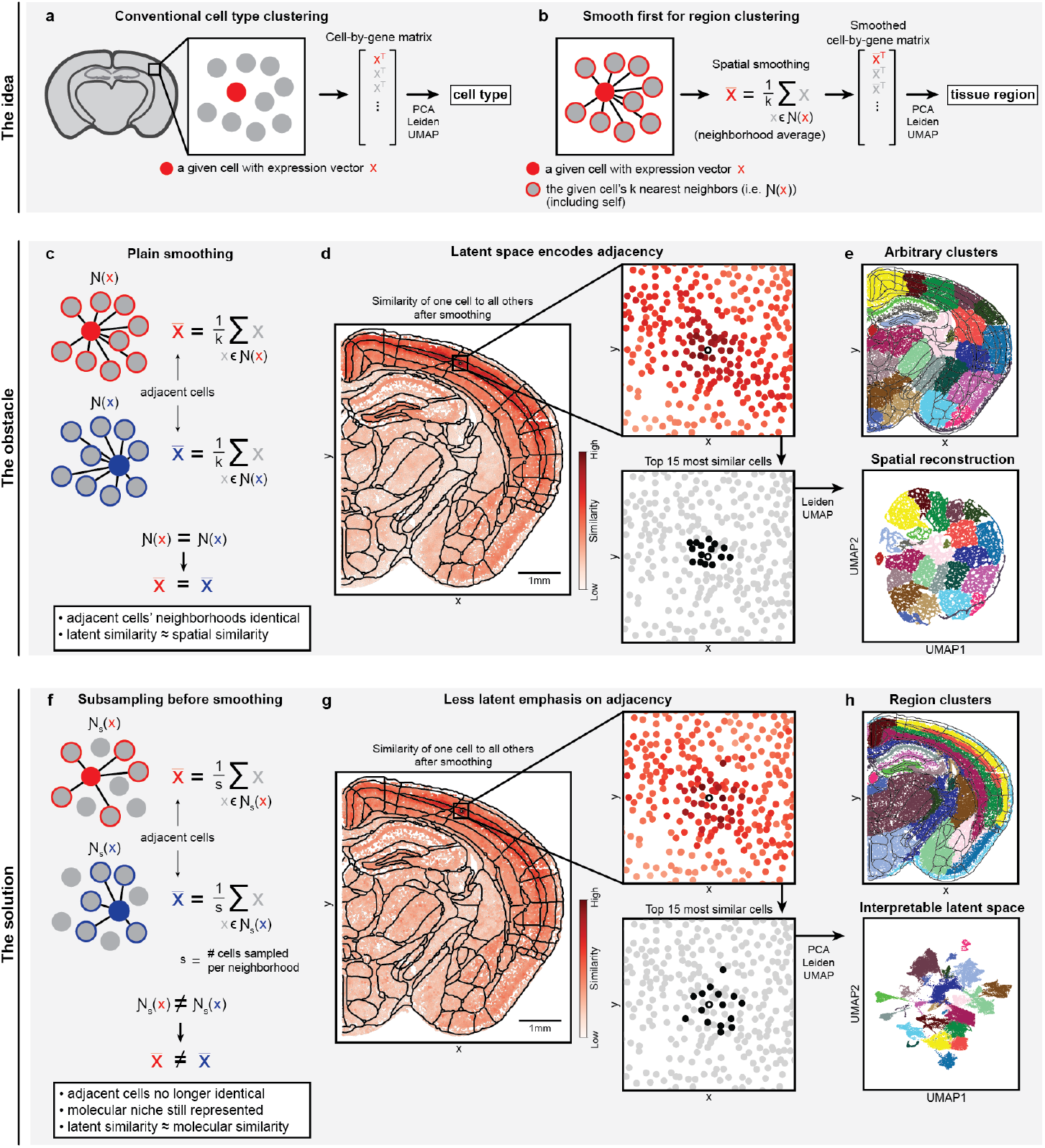
Schematic of the smoothing idea, autocorrelation obstacle, and subsampling solution. **a)** Schematic of conventional cell type clustering. Each dot represents a cell. **b)** Schematic of recent region clustering methods, in which expression features are smoothed before applying conventional single-cell clustering tools. The dots shown are the same as those in a). **c)** Schematic demonstrating the main limitation of smoothing. The dots shown are the same as those in a) and b). The red and blue filled dots represent physically adjacent cells. **d)** Results of comparing the smoothed representation of one cell (represented as a dot with white fill and black outline) to all others in the tissue. Similarity is defined as the dot product between normalized expression feature vectors. An anatomical wireframe from the Allen Brain Atlas is overlaid for comparison with conventional mouse brain anatomy^16^. **e)** Results of conventional single-cell clustering on smoothed data. PCA was omitted to emphasize physical reconstruction. For results that include PCA, see Fig. 2a. **f)** Schematic demonstrating a simple solution to the problem shown in c). **g-h)** Same as d-e) but with random neighborhood subsampling and including PCA during clustering. For e,h), Leiden resolutions were set such that clustering yielded 23 clusters.

We demonstrated this phenomenon in a STARmap PLUS dataset of mouse brain from a recent atlas^15^. We began by smoothing over neighborhoods of size *k*=30 and then visualizing the similarity of one neocortical cell’s smoothed expression vector to all other cells in the dataset. While the given cell was appropriately less similar to subcortical cells and more similar to other neocortical cells, it was indeed the most similar to its immediate spatial neighbors (Fig. 1d). As a result, after applying Leiden and UMAP, the latent space appeared to reconstruct the physical space, generating spatially contiguous yet arbitrary cluster labels (Fig. 1e; for an example of equivalent results that include PCA, see Fig. 2a). This can also be viewed as a combined consequence of randomness of neighborhood cell-type compositions and noisiness of single-cell expression, making the smoothed features of neighboring cells uniquely similar. Indeed, when expression features are randomly shuffled across the tissue, this phenomenon persists and is more pronounced, with the tissue again being physically reconstructed in latent space (Supplementary Fig. 1). While PCA can deemphasize the effects of spatial autocorrelation, we find that it is not sufficient to eliminate them, necessitating a more comprehensive solution (Supplementary Fig. 2).

**Figure 2.**
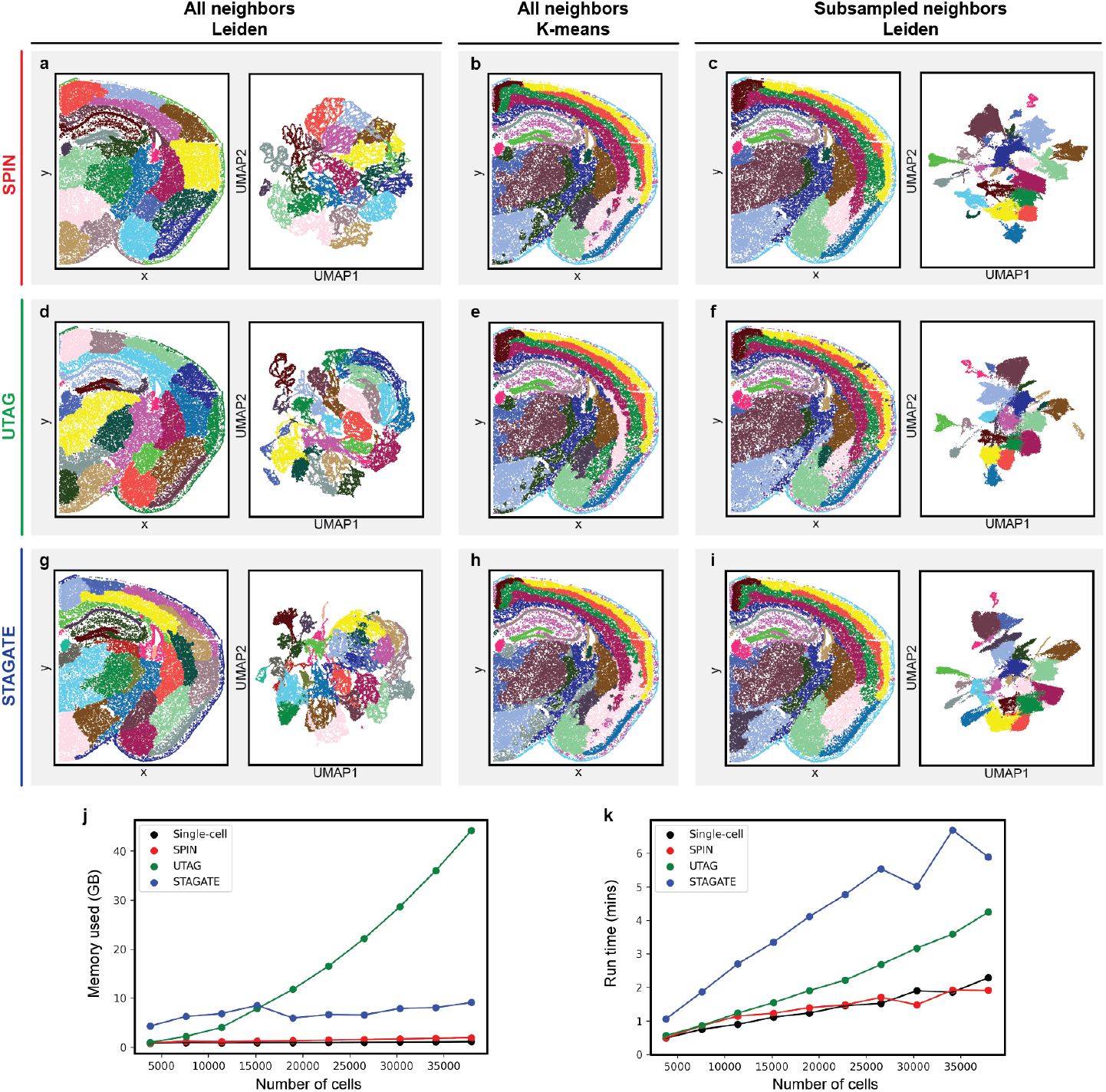
Comparing SPIN to existing methods. **a)** Leiden clustering results after smoothing (without subsampling). Same as Fig. 1e but including PCA with 50 PCs. **b)** K-means clustering results after smoothing **c)** Leiden clustering results after neighborhood subsampling followed by smoothing, i.e. SPIN. Same as Fig. 1h. **d)** Leiden clustering results using UTAG. **e)** K-means clustering results using UTAG. **f)** Leiden clustering results from subsampling followed by UTAG. **g)** Leiden clustering results using STAGATE. **h)** K-means clustering results using STAGATE. **i)** Leiden clustering results from subsampling followed by STAGATE. **j)** Memory usage of each method applied to different amounts of cells. **k)** Run time of each method applied to different amounts of cells. “Single-cell” in j,k) refers to the analysis shown in Fig. 1a. Curves were created by running each method on subsets of the mouse brain dataset shown in a-i), from 10% to 100% of the total number of cells, in increments of 10%. Subsampling is included as part of each method. For a-i), k-means and Leiden were performed such that clustering yielded 23 clusters.

We identified a simple solution to this issue in which each cell’s spatial neighborhood is randomly subsampled before smoothing (Fig. 1f). We reasoned that, given the number of samples is sufficiently large to represent each cell’s molecular niche, subsampling would allow adjacent cells to differ in their *exact* neighbor compositions while still capturing their *general* molecular compositions. We identified neighborhoods of size *k*=30 cells, randomly subsampled each neighborhood with samples of size *s*=10 cells, and then repeated smoothing. Indeed, when comparing one cell to all others as performed above, the subsampling approach placed less emphasis on spatial adjacency (Fig. 1g). As a result, clustering identified the major molecular tissue regions expected in the dataset, including the neocortical layers as well as many subcortical structures (Fig. 1h; for region labels, see Fig. 3d). Furthermore, the latent space appeared to capture features of anatomical structure, as evidenced by the one-dimensional manifold corresponding to the laminar structure of the neocortex. These exact values of *k* and *s* were determined by manual variation and inspection of the results. While more extensive optimization may be possible, we find that these parameters are sufficient to identify major anatomical regions in most of the SRT datasets explored here. Finally, despite the randomness of subsampling, we find that multiple runs with different random seeds show consistent results, although potentially requiring different clustering resolutions to identify the same set of regions (Supplementary Fig. 3).

**Figure 3.**
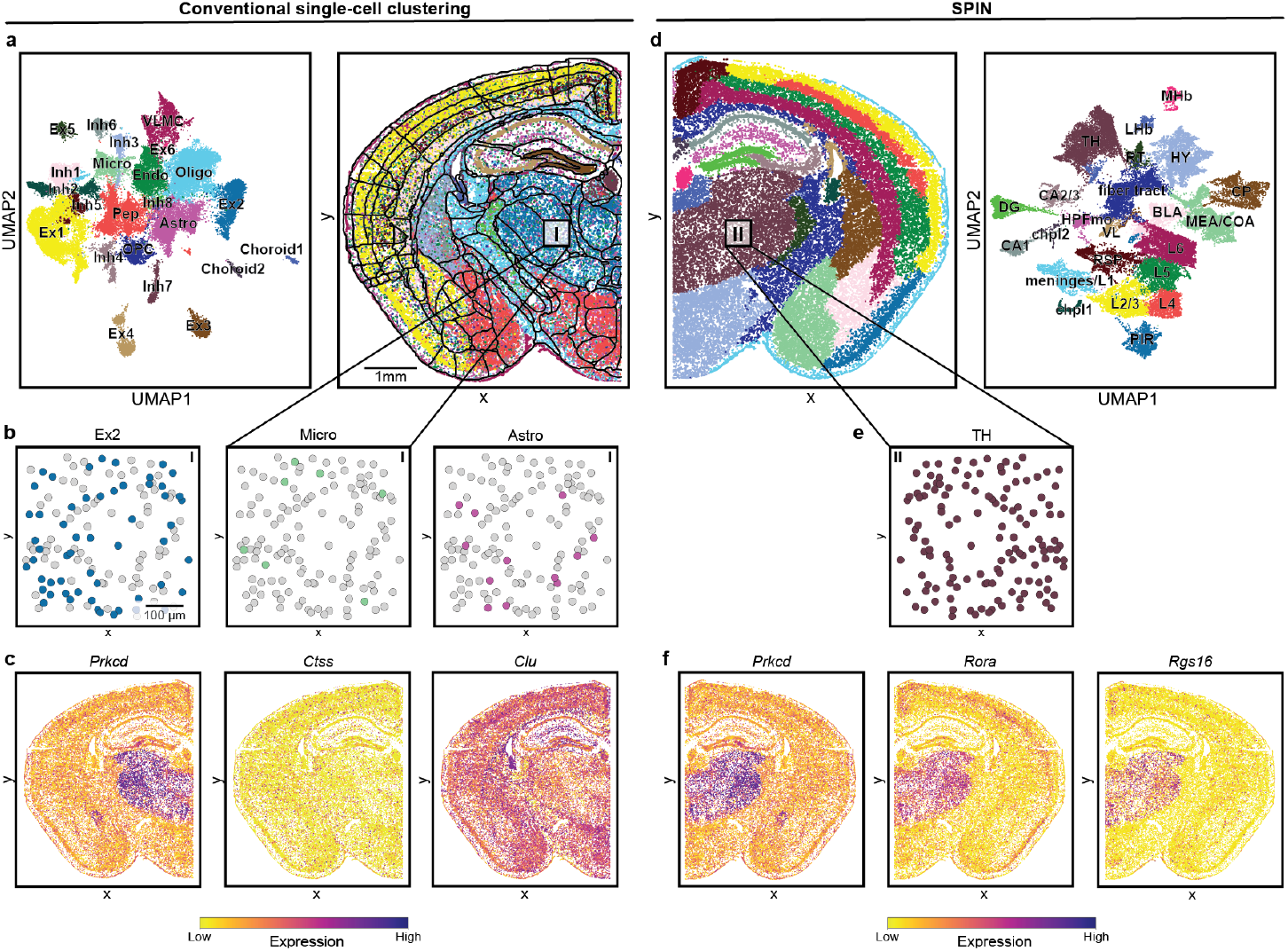
Spatial DEG analysis with SPIN. **a)** Visualization of cells in UMAP space colored and labeled by molecular cell type (left). Visualization of cells in UMAP space (left) and *in situ* (right) colored by cell type. An anatomical wireframe from the Allen Brain Atlas is overlaid for comparison of cell type distributions with conventional anatomy^16^. Inset box I indicates area of zoom-in view in b). **b)** Examples of cell type distributions within the Thalamus (TH). **c)** Marker genes for each cell type directly above in b). **d)** Visualization of cells *in situ* (left) and in UMAP space (right) colored by molecular tissue regions, which are generated by subsampling and smoothing the data before clustering. Same as Fig. 1h and Fig. 2c. Same dataset as in a) but horizontally reflected. Inset box II indicates area of zoom-in view in e). Visualization of labeled regions in UMAP space colored and labeled by region (right). **e)** Cells from the same Thalamic region as b). **f)** Visualization of the top three marker genes *in situ* for the Thalamic cluster in e) (left to right). For a,d), Leiden resolutions were set such that each approach yielded 23 clusters.

Several alternative approaches to subsampling can also be derived based on this principle. For example, an alternative aggregation approach is to concatenate each neighbor-hood’s gene expression profiles rather than average them. Single-cell gene expression vectors of a cell and its spatial nearest neighbors (without subsampling) can be concatenated in order of distance, from nearest to furthest to the given cell, forming a neighborhood vector of length *dk* where *d* is the number of genes measured and *k* is the number of neighbors (Supplementary Fig. 4a). This concatenation approach assigns an ordering to each cell’s neighborhood and relies on the fact that even identical neighborhoods will have different orderings; as each cell is closest to itself, the order of each cell’s neighborhood vector will always begin with itself and thus differ from all other cells’. Furthermore, even if adjacent cells have identical neighborhoods, their relative distances to each neighbor likely differ, leading to additional variation in their concatenation orderings. Clustering on concatenated features yields largely the same molecular tissue regions and latent representation as the subsampling approach (Supplementary Fig. 4b,c). While one may wonder whether the distance-based ordering encodes relevant biological information, we find that randomly shuffling the ordering of each cell’s neighborhood produces nearly identical clustering results, suggesting that the *absolute* ordering is irrelevant so long as the *relative* ordering of adjacent cells is different (Supplementary Fig. 4d,e). However, as concatenation increases the size of the data matrix by a factor of *k*, the PCA step is much more computationally costly than conventional single-cell analysis. On the other hand, the subsampling approach is computationally equivalent to single-cell analysis, apart from the overhead required for smoothing.

Finally, rather than modifying the smoothing strategy, one can instead vary the clustering strategy. For instance, k-means clustering can be applied to plainly smoothed (without subsampling) data to recover largely the same tissue regions as Leiden with subsampling or concatenation (Supplementary Fig. 5a). This is due to the fact that k-means does not depend on pairwise similarities between cells in order to determine clusters. Rather, a fixed number of cluster centroids are initialized, and cells are iteratively assigned to each centroid without individual comparisons. Alternatively, one can avoid clustering altogether by utilizing clusterfree methods to identify molecular tissue regions. For instance, non-negative matrix factorization (NMF)^17^ can be applied to plainly smoothed data to calculate factors corresponding to distinct spatial expression patterns (Supplementary Fig. 5b). This approach is similar to k-means in that it also does not rely on individual comparisons. However, NMF does not generate categorical cluster labels which may be necessary for downstream analysis. Furthermore, neither k-means nor NMF produce latent spaces that are compatible with downstream analyses relying on nearest-neighbor graphs, such as Leiden, UMAP, or trajectory inference. This is because k-means does not explicitly calculate a latent embedding, while the latent representation given by NMF still embeds physically adjacent cells closest to one another, again causing latent nearest neighbors to reflect spatial proximity. Thus, among the strategies explored above, we find subsampling to be the most efficient and convenient solution.

An important consideration when using the above approaches is that molecular regions may exist on varying length scales^1^. The number of neighbors used for smoothing, *k*, determines which length scale the spatial expression patterns are detected on. For instance, smoothing over a large *k* might allow identification of large regions such as cortical layers while ignoring smaller-scale structures such as meninges or vasculature, and vice versa. We demonstrated this by varying *k* and comparing the resulting molecular region clusters (Supplementary Fig. 6a). Between *k*=1 and *k*=5, clustering identified small regions corresponding to vasculature (Supplementary Fig. 6b,c). Marker genes indicated these regions were indeed composed of endothelial cells and pericytes (Supplementary Fig. 6d). However, at *k*=30, the same vascular cells were instead classified based on the cortical layers they occupied, with top marker genes corresponding to the given cortical layer (Supplementary Fig. 6e).

Altogether, we found that smoothing SRT data increases spatial autocorrelation and causes latent reconstruction of physical space, obscuring the underlying spatial molecular patterns. However, several approaches outlined above can mitigate this issue, with subsampling in particular offering efficiency and complete compatibility with the conventional single-cell toolkit. We refer to this subsampling and smoothing approach as <u>sp</u>atial <u>in</u>tegration (SPIN).

### Mitigating spatial autocorrelation in existing methods

Because existing methods for molecular region clustering in SRT data can be expressed as more complex variations on smoothing, we sought to determine whether they also cause artifactual spatial reconstruction of the data. We began by smoothing (without subsampling) and clustering as shown above for comparison (Fig. 2a). Here we did not omit PCA, so the physical reconstruction in UMAP space was less pronounced. However, it was still evident based on the quilting of the arbitrary, spatially contiguous tiles in latent space. We then plotted the prior results from k-means and SPIN to visualize the expected molecular regions (Fig. 2b,c). For our first comparison, we applied UTAG to the same mouse brain dataset, which yielded similar results to smoothing without subsampling (Fig. 2d). While one could argue that this may be due to suboptimal parameter selection, we found that UTAG with k-means identified the expected molecular regions, indicating that the underlying spatial expression features were present in the latent representation despite being obscured by Leiden and UMAP (Fig. 2e). Indeed, when a randomly subsampled adjacency matrix was provided as input, the expected regions were detectable using Leiden and UMAP (Fig. 2f). This suggests that UTAG also increases spatial autocorrelation which can be mitigated by subsampling. We then determined whether this issue affects STAGATE’s more complex graph neural network-based model. Again, applying Leiden and UMAP to the latent space output produced spatially contiguous yet arbitrary clusters (Fig. 2g), while k-means identified the expected regions (Fig. 2h), and inputting a subsampled adjacency matrix enabled Leiden and UMAP to also detect the expected regions (Fig. 2i). Thus, we find that spatial autocorrelation is increased and can be mitigated by subsampling even in more complex graph neural network-based models such as STAGATE.

Given that subsampling enabled each method to identify comparable molecular tissue regions, we next sought to compare the computational efficiency of each method. To do so, the conventional single-cell approach, SPIN, UTAG, and STAGATE were each used to cluster subsets of the same STARmap PLUS mouse brain dataset, from 10% of the data to 100%, in increments of 10%. As the majority of these methods do not rely on GPU acceleration, we chose to compare them using CPU only. SPIN showed the least memory usage and was comparable to single-cell clustering, while STA-GATE showed consistently higher usage across all amounts of cells, and UTAG usage appeared to exponentially increase with more cells (Fig. 2j). With respect to run time, SPIN again appeared equivalent to single-cell clustering, while UTAG and STAGATE became less efficient as the number of cells increased (Fig. 2k). Thus, considering both memory usage and run time, we find that SPIN is more efficient than the existing region identification methods shown here.

For the remainder of this work, we further demonstrate the power and convenience of SPIN by leveraging additional conventional single-cell tools to jointly analyze various SRT datasets.

### DEG analysis identifies region marker genes

Cell type clustering is typically followed by DEG analysis for molecular characterization of each cell type. We demonstrated this process by applying conventional cell type clustering and DEG analysis to the same STARmap PLUS sample of mouse brain shown above (Fig. 3a). Clusters were assigned cell type labels based on their respective marker genes. As expected, conventional anatomical regions contained mixtures of cell types. For instance, the thalamus contained a mixture of glutamatergic neurons (Ex2), microglia (Micro), and astrocytes (Astro) (Fig. 3b). The marker genes defining these cell types demonstrated various extents of spatial patterning. The glutamatergic neuron marker *Prkcd* appeared to be spatially distributed uniquely within the thalamus, while the microglial marker *Ctss* appeared uniformly distributed, and the astrocyte marker *Clu* appeared spatially patterned in some areas such as the cortex yet uniform in others (Fig. 3c).

On the other hand, clustering on subsampled and smoothed data yielded spatially contiguous molecular region labels, identifying marker genes specific to each region (Fig. 3d,e). For example, the top three marker genes for the thalamus region all appeared to be spatially variable genes (SVGs) expressed within the thalamus (Fig. 3f). This suggests that smoothing successfully removes sparse, non-spatial gene expression patterns, instead maintaining and propagating patterns of SVGs. Thus, when combined with SPIN, DEG analysis is capable of identifying SVG markers for each molecular tissue region.

### Integration aligns spatial transcriptomic features across species

Given that SPIN combined with clustering and DEG analysis identified molecular tissue regions and their corresponding marker SVGs, we wondered whether single-cell integration methods could further enable joint characterization of regions across multiple datasets. We reasoned that the *local* operation of smoothing should allow integration to align spatial expression features across datasets with arbitrary *global* morphologies. In other words, while differences in morphology between samples can prevent accurate physical registration, molecular comparison of *local* cellular neighborhoods is independent of *global* morphology, allowing alignment of samples with arbitrary shapes. We chose to demonstrate this by comparing a STARmap PLUS dataset of mouse frontal cortex with a STARmap dataset of marmoset frontal cortex, the former from a recent mouse brain atlas^15^ and the latter newly collected for this study. We first identified cell types for reference by performing single-cell integration using Harmony^18^ followed by two levels of clustering as described above (Supplementary Fig. 7). Major excitatory neuron, interneuron, and non-neuronal cell types were detected in the data (Fig. 4a-c). We then performed subsampling and smoothing on each dataset independently with *k*=30 and *s*=12, followed by the same integration and clustering approach. Region clusters were then annotated by referencing mouse and marmoset anatomical atlases^15,19–21^. Clustering revealed the laminar neocortical structure in each dataset (Fig. 4d,e), which was reflected in the one-dimensional shape of the UMAP embedding (Fig. 4f). Integration appeared to successfully align spatial expression features from each species, as indicated by the overlap between latent representations of cells from each dataset (Fig. 4f, inset). Notably, integration appeared to preserve and identify real dataset-specific regional differences, as demonstrated by the lack of species overlap in the anterior commissure (aco), piriform area (PIR), and taenia tecta (TT) regions, which were present in the mouse sample but absent in the marmoset sample.

**Figure 4.**
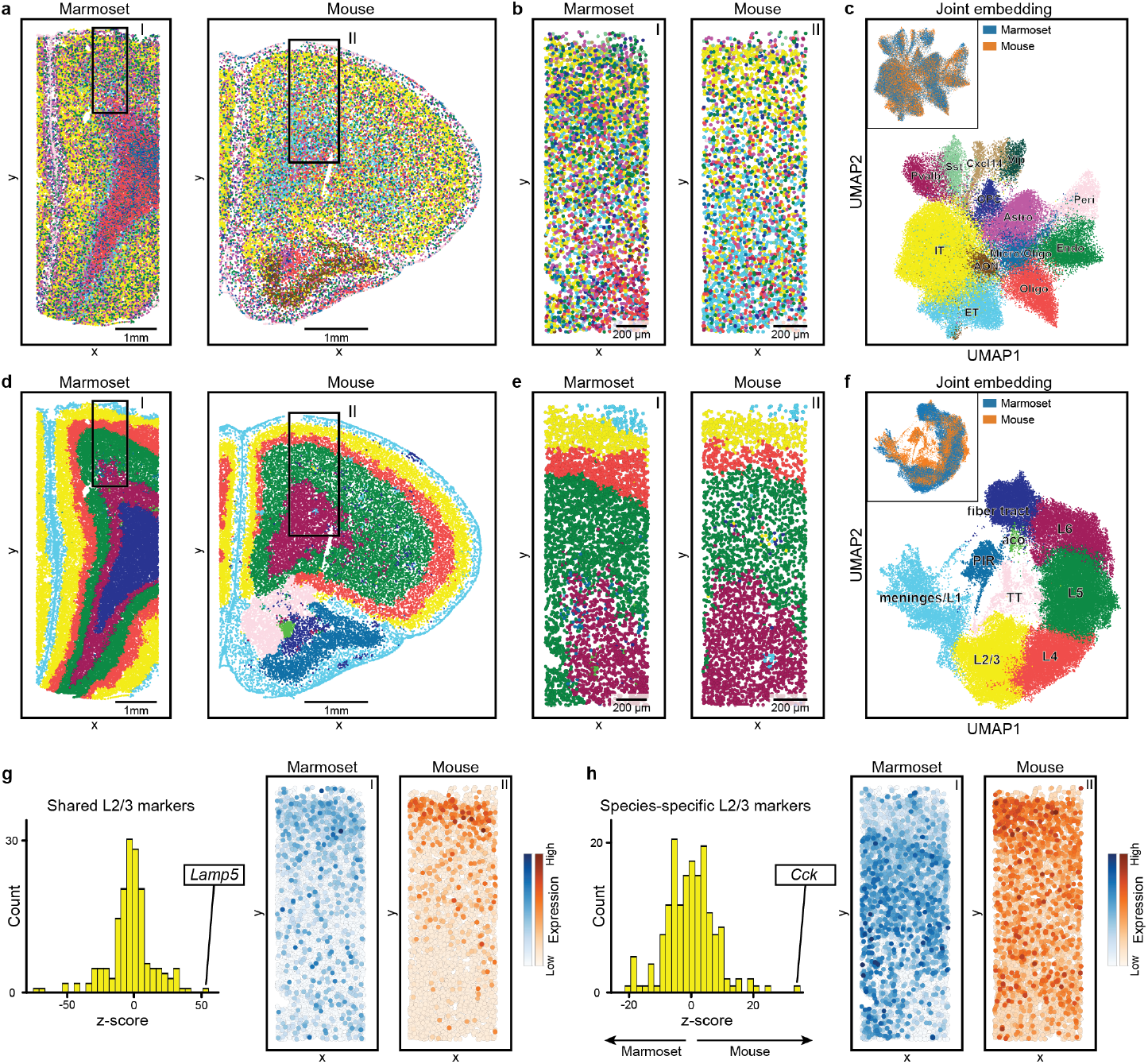
Spatial integration across species with SPIN. **a)** Visualization of tissue *in situ* colored by molecular cell type. The marmoset STARmap dataset contained measurements of 454 genes across 42,458 cells, and the mouse STARmap PLUS dataset contained measurements of 1,022 genes across 29,623 cells. After integration, each dataset contained measurements across 174 genes (the genes measured in both datasets). Inset boxes I and II indicate areas for zoom-in views used in b), e), and g), and h). **b)** Zoom-in views of the tissue colored by cell type. **c)** Visualization of tissue in UMAP space colored and labeled by cell type. Inset shows tissue in UMAP space colored by species. **d-f)** Same as a-c) but labeled by molecular tissue regions generated by SPIN (i.e. clustering the subsampled and smoothed data). **g)** Histogram of L2/3 marker scores for each gene (left). Scores correspond to t-test results for a given gene across cells in L2/3 versus cells in all other regions (i.e. Scanpy’s *sc*.*tl*.*rank_genes_groups*). The top joint marker for L2/3 in both species was *Lamp5*, which is visualized *in situ* (right). **h)** Histogram of species-specific L2/3 marker scores for each gene. Scores were calculated as in g) but by comparing all L2/3 cells based on species. The top species-specific marker was *Cck*, which is visualized *in situ* (right).

Given the ability to molecularly align regions across datasets, we further sought to calculate shared and species-specific spatial transcriptomic features. Shared marker genes for each region were calculated using DEG analysis across region groups using data from both species, yielding region marker genes that were shared by both species. For instance, the top shared marker for L2/3 was *Lamp5*, a canonical marker for that region, which was confirmed by *in situ* visualization of STARmap measurements (Fig. 4g; see Supplementary Fig. 8a-c for plots of the full tissue). We then calculated species-specific region markers by isolating cells within a given region and performing DEG analysis across species. *Cck* was identified as the top differentially distributed gene (DDG) in the L2/3 region, which was also visually evident *in situ* (Fig. 4h, Supplementary Fig. 8d-f). Notably, *Cck* is expressed in both excitatory and inhibitory neurons in mice^22^, indicating that the difference in spatial distributions between species is not due to an underlying difference in a single cell type. Thus, this approach is capable of identifying differences in spatial distributions that are independent of cell types. Furthermore, *Cck* neurons have been shown to regulate opiate antagonism^23,24^, satiety signaling^25^, and learning and memory^26^, indicating that this difference in spatial distribution between species may have functional implications. Finally, we visualized the top marmoset specific L2/3 marker, *Pde1a*. However, just as the magnitude of its z-score was lower than that of *Cck*, the difference between species was more difficult to visualize by eye (Supplementary Fig. 8g-i).

### Integration aligns spatial transcriptomic features across technologies

While single-cell integration methods are often used to identify differences across datasets, they are also used to merge datasets collected using diverse experimental technologies^27^. This has enabled the creation of single-cell atlases that leverage the unique strengths of multiple technologies, such as single-cell and single-nucleus RNA sequencing^28,29^. SRT technologies also vary in their strengths, with methods such as Visium^30^ measuring more genes with lower spatial resolution and methods such as STARmap measuring fewer genes with higher spatial resolution. As SRT atlases are increasingly being generated using a variety of technologies^6,15,31–35^, we sought to test whether integrating smoothed data facilitates joint characterization of spatial datasets from different technologies.

We thus used SPIN to spatially integrate three open-source SRT datasets of mouse brain from technologies with decreasing spatial and increasing transcriptomic resolutions: STARmap PLUS, Slide-seqV2^36^, and Visium. To accommodate their different spatial resolutions, smoothing was performed with varying neighborhood sizes across each dataset (Methods). Integration and clustering yielded molecular tissue regions that reflected the underlying anatomy within each dataset, despite broad differences in spatial resolution, orientation, and field of view (Fig. 5a). Regions represented in UMAP space appeared well-aligned across technologies as evidenced by dataset mixing (Fig. 5b). Notably, regions were accurately identified despite the limited field of view of the Slide-seqV2 sample as well as the more posterior anatomical position of the Visium sample. Finally, an additional layer 6b region (L6b) was detected, which was not found in the STARmap data alone.

**Figure 5.**
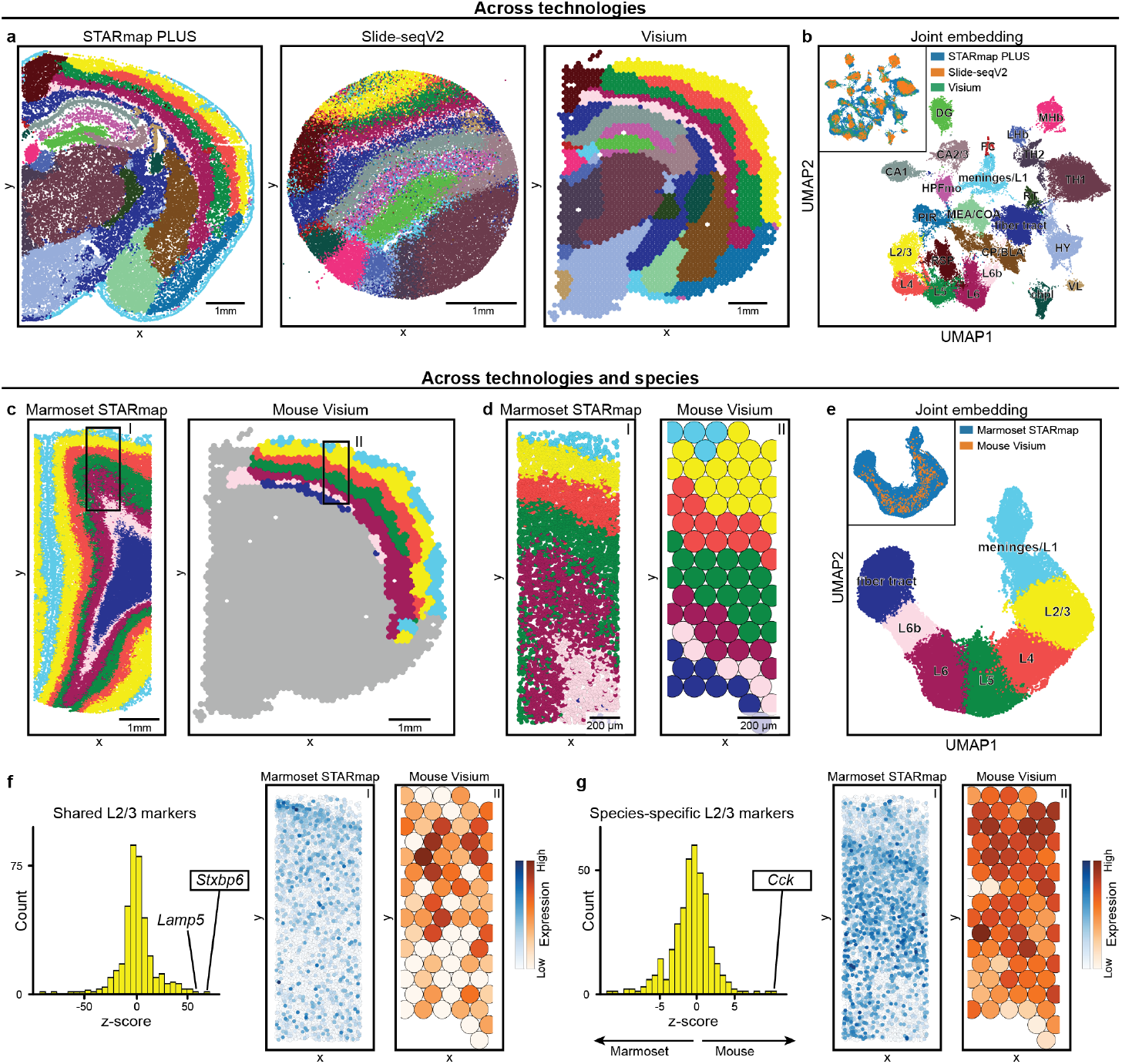
Spatial integration across experimental technologies with SPIN. **a)** Visualization of tissue from each technology *in situ* colored by molecular tissue region. From left to right, technologies decrease in spatial resolution and increase in transcriptomic resolution. The STARmap PLUS dataset contained 1,022 genes measured across 37,928 cells, the Slide-seqV2 dataset contained 4,000 genes measured across 41,786 spots, and the Visium dataset contained 18,078 genes measured across 2,688 spots. After integration, each dataset contained measurements across 521 genes. **b)** Visualization of dataset in UMAP space colored and labeled by molecular tissue region. Inset is the same but colored by technology. **c)** Visualization of tissue *in situ* colored by molecular tissue region. After integration, each dataset contained measurements across 401 genes. Inset boxes I and II indicate areas for zoom-in views used in d), f), and g). **d)** Zoom-in views of the tissue colored by region. **e)** Visualization of cells/spots in UMAP space colored and labeled by molecular tissue region. Inset shows cells/spots in UMAP space colored by species. **f)** Histogram of L2/3 marker scores for each gene (left). The top ranked marker for L2/3 in both species was *Stxbp6*, which is visualized *in situ* (right). **g)** Histogram of species-specific L2/3 marker scores for each gene. The top species-specific marker was *Cck*, which is visualized *in situ* (right).

Beyond the creation of consensus atlases, we reasoned that cross-technology spatial integration could enable comparison across datasets that differ both in technology and in other properties. For instance, consider the scenario where one wants to compare molecular regions across marmoset and mouse neocortex yet only has access to marmoset data collected using one technology and mouse data collected using a different technology. In this case, the ability to spatially integrate across both species and technology would be necessary. To test whether this was possible, we isolated the above neocortical regions from the Visium mouse brain dataset and performed SPIN with the prior marmoset neocortex STARmap data as shown above. All cortical, fiber tract, and meninges regions from the prior cross-species integration were once again detected, and the latent representation appeared similar as well (Fig. 5c-e). Furthermore, integration again allowed the identification of an additional L6b region that was not detected in the STARmap data alone. Because the mouse Visium dataset measured more genes than in the prior mouse STARmap PLUS dataset, the overlap with the marmoset data was greater, allowing the detection of additional genes, such as *Stxbp6*, that were enriched in L2/3 (Fig. 5f). However, *Cck* remained the top species-specific gene in L2/3 (Fig. 5g), matching the results from our previous cross-species analysis that only used STARmap data.

### Trajectory inferences identifies continuous molecularly defined spatial axes

While SPIN allowed joint characterization of discrete tissue regions, the one-dimensional latent structure of the neocortical data suggested the presence of meaningful continuous variation as well (Fig. 3d, Fig. 4f, Fig. 5b,e). The conventional single-cell tools used to characterize such molecular gradients are referred to as trajectory inference or pseudo-time methods^37^. In the context of single-cell analysis, the gradients identified by these methods are typically interpreted as developmental trajectories. We reasoned that, when applied to smoothed data, these same methods would instead identify spatial molecular gradients within the tissue. We thus applied diffusion maps^38^, a popular trajectory inference method, to the prior smoothed and integrated marmoset and mouse neocortical STARmap datasets. As the neocortex is known to be organized into layers L2/3-L6 ordered by depth,we isolated cells from these cortical region clusters for demonstration. Indeed, the first diffusion component (DC1) formed a one-dimensional measure of depth along the neocortex (Fig. 6a). Notably, this measure was not physically uniform in physical space; for example, a large change in DC1 occurred within a small physical portion of the tissue corresponding to L4. Accordingly, we termed this measure “molecular depth”. Importantly, calculation of molecular depth is unsupervised, independent of cell type clustering, and can be performed across multiple datasets, unlike previous supervised, cell type-based approaches^30,39^.

**Figure 6.**
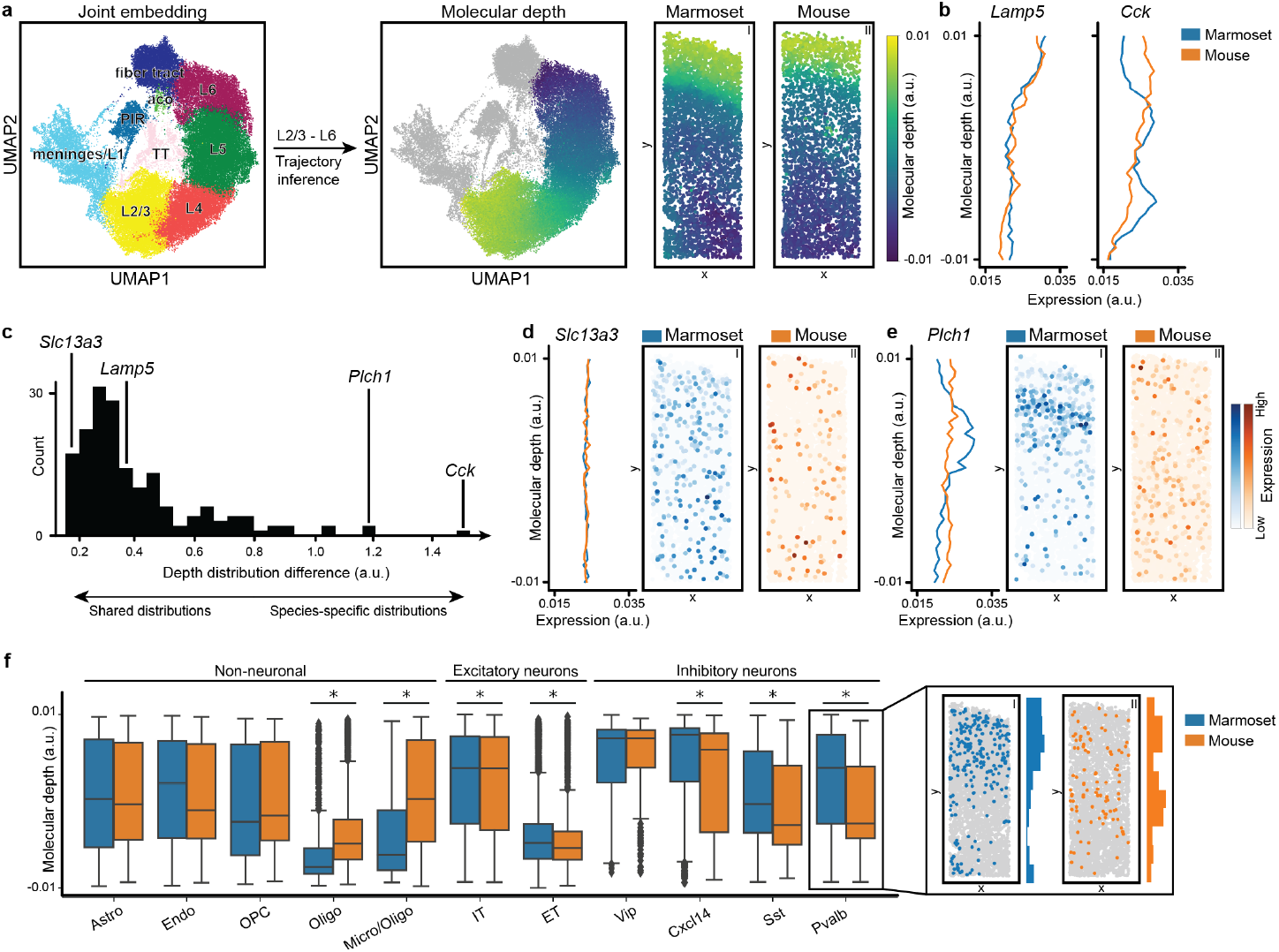
Joint spatial trajectory inference across species with SPIN. **a)** Visualization of tissue in UMAP space colored and labeled by molecular tissue region (left; same as in Fig. 4f). Visualization of diffusion component 1 (DC1), which we termed “molecular depth”. Molecular depth visualized over the tissue in UMAP space (middle). Zoom-ins of the same brain tissue regions as in Fig. 4b,e,g,h, colored by molecular depth (right). **b)** Depth distributions for the top shared and species-specific L2/3 genes from the analysis shown in Fig. 4g,h. Cells were binned according to depth, and expression was averaged across cells within each bin to calculate a depth distribution for each gene. **c)** A histogram of depth distribution differences for each gene across species. **d)** Depth distribution plots and *in situ* visualization of *Slc13a3*, which had the lowest depth distribution difference. **e)** Depth distribution plots and *in situ* visualization of *Plch1*, which had the second highest depth distribution difference. **f)** Boxplot showing the depth distributions of cells from each cell type and each species. Boxes extend from the first to the third quartile with the middle line showing the mean. Whiskers extend above and below the first and third quartiles by 1.5 times the interquartile range. Points outside this range are considered outliers and depicted as individual diamonds. For each cell type, distributions were compared across species using a two-sided Mann-Whitney U-test and \**P*<0.01 after Bonferroni correction for multiple comparisons. Zoom-in views of the *in situ* distributions of Pvalb interneurons are shown for illustration (right). Paired histograms correspond to the physical depths of the cells along the y-axis of the tissue for comparison to molecular depth.

We leveraged this approach to identify genes with differing neocortical molecular depth across species. For each gene, a depth distribution was calculated by binning cells according to molecular depth and calculating the average expression of the gene within each bin. To compare a given gene’s depth distribution across species, we summed the absolute values of the differences in expression between species at each depth bin. To validate this metric, we first plotted the depth distributions of *Lamp5* and *Cck*, which recapitulated the spatial patterns found during our previous DEG analysis (Fig. 6b). We then calculated the depth distribution differences between species for all measured genes (Fig. 6c). Indeed, *Lamp5* showed a relatively low difference across species, whereas *Cck* showed the highest difference. The lowest difference was in *Slc13a3*, which appeared spatially uniform in both species (Fig. 6d). The second highest difference was in *Plch1*, which appeared spatially uniform in mouse neocortex yet uniquely expressed in L4 in marmoset (Fig. 6e).

We then compared the molecular depth of cell types across species. For a given cell type, depth distributions for each species were compared. Significant differences in depth were observed within oligodendrocytes (Oligo, Micro/Oligo), excitatory neurons (IT, ET), and Cxcl14, Sst, and Pvalb interneurons (Fig. 6f, left). Out of these significant groups, modest differences were found between excitatory neurons, which were more superficial in marmoset neocortex compared to mouse. Larger differences were found between oligodendrocytes, which were much deeper in marmoset neocortex. Finally, Cxcl14, Sst, and Pvalb interneurons appeared much more superficial in the marmoset neocortex compared to mouse, recapitulating results from prior supervised analyses^40,41^. To visualize an example of the difference in cell type molecular depths, we plotted Pvalb interneurons *in situ* in both species along with physical depth histograms based on each sample’s y-axis (Fig. 6f, right). The *in-situ* visualization and physical depth distributions both appeared to recapitulate the molecular depth distributions and their differences between species.

## Discussion

We demonstrated that multiple smoothing-based methods physically reconstruct SRT data in latent space, preventing accurate characterization of spatial transcriptomic features. We introduced several simple modifications that circumvent this issue, including a subsampling approach that can improve the performance of existing methods as well as form the basis of a new minimal method for spatial characterization that we call SPIN. We then showed that SPIN extends the utility of conventional single-cell tools to SRT data, creating spatial analogies of each tool. This allowed the unsupervised identification of genes whose spatial distributions differ across species. It also enabled the unsupervised identification of genes and cell types that differ in neocortical molecular depth across species. Altogether, we introduced a simple, efficient, and effective approach to analyzing spatial transcriptomic features using conventional single-cell tools.

Drawbacks of the smoothing and clustering strategies mentioned here may arise at small or large length scales. In the case of small length scales, the subsampling approach may fail due to the inability to capture the relevant regional features within so few cells. For example, smoothing with *k*=5 would require using *s*<5 subsampled cells, which may not be sufficient to capture the molecular diversity of the regions of interest. However, in this case, one may benefit from the concatenation or k-means approaches to avoid subsampling altogether. On the other hand, large length scales may prohibit the use of concatenation due to computational inefficiency. Furthermore, the requirement of choosing a single *k* prevents the identification of molecular regions along both small and large length scales simultaneously. However, one could likely utilize the hierarchical subclustering approach often used in single-cell clustering to iteratively identify large-scale regions, their constituent smaller-scale subregions, and so on.

We anticipate SPIN will be utilized for two primary applications. First, given the efficiency of spatial smoothing as well as conventional single-cell analysis methods, we expect the use of SPIN for combining and comparing spatial transcriptomic features across emerging large-scale SRT atlases^6,15,31–35^. As anatomy corresponds with function, such atlas-level spatial comparisons may be able to identify transcriptional underpinnings of functional differences between species or conditions. Second, we anticipate that SPIN will be used to generate spatial analogies for several additional classes of single-cell methods not explored here. For instance, SPIN may be capable of generalizing single-cell multimodal integration methods to spatial pattern alignment across multiple modalities. On the other hand, novel integration methods such as SATURN^42^ allow flexible alignment of cells across species based on transcriptional features as well as protein embeddings of the corresponding genes, yielding cell embeddings and clusters that are more relevant to biological function. Thus, when combined with SPIN, they may also enable more functionally informative characterization of molecular tissue regions across species. Furthermore, cluster-free differential abundance methods such as Milo^43^ and CNA^44^ allow identification of cell type populations that are more abundant in a given dataset compared to others. Thus, such methods applied alongside SPIN may be capable of revealing differentially abundant molecular tissue regions across datasets. Finally, novel cluster-free DEG methods such as LEMUR^45^ and miloDE^46^ could be paired with SPIN to calculate DDGs in SRT data, potentially allowing flexible, cluster-free identification of spatial transcriptomic differences between tissues.

## Data availability

The marmoset brain STARmap dataset is available at https://zenodo.org/record/8092024. A mouse STARmap PLUS dataset is included for demonstration of SPIN.

## Code availability

Source code, usage principles, and a tutorial notebook for SPIN are available at https://github.com/wanglab-broad/spin.

## Acknowledgements

We thank A. Nadig, T. Kamath, and D. Barabasi for helpful conversations and comments on the manuscript. X.W. acknowledges the support from Searle Scholars Program, Thomas D. and Virginia W. Cabot Professorship, and Edward Scolnick Professorship. This paper was typeset with the bioRxiv word template by @Chrelli: www.github.com/chrelli/bioRxiv-word-template.

## Author contributions

K.M. conceived of the method. K.M. and X.W. designed the research. K.M. and M.W. performed data analysis and prepared the manuscript. Q.Z. collected the marmoset tissue. Y.Z. performed STARmap on the marmoset sample and collected the data. J.H. preprocessed the marmoset imaging data. K.M., M.W., Y.Z., J.H., and X.W. critically revised the manuscript. X.W. supervised the study.

## Materials and Methods

### Marmoset sample collection

All marmosets were either family group-housed or pair-housed in large cages with a variety of perches and enrichment devices. All marmoset cages are in spacious rooms with environmental control of temperature (23– 28°C), humidity (40–72%), and 12 hr light/dark cycle. All marmosets received regular health checks and behavioral assessment from MIT DCM veterinary staff and researchers. All animal procedures were conducted with prior approval by the MIT Committee for Animal Care (CAC) and following veterinary guidelines.

Adult marmosets (2.5 years old, female) were deeply sedated by intra-muscular injection of ketamine (20–40 mg kg −1) or alfaxalone (5–10 mg kg −1), followed by intravenous injection of sodium pentobarbital (10–30 mg kg −1). When the pedal withdrawal reflex was eliminated and/or the respiratory rate was diminished, animals were transcardially perfused with ice-cold sucrose-HEPES buffer. Whole brains were rapidly extracted into fresh buffer on ice. Brain regions were dissected using a marmoset atlas as reference^47^ and were snap-frozen in liquid nitrogen.

### Gene panel selection

Marker genes and most differentially expressed genes were extracted from single-cell RNA-sequencing studies^40,48^ that surveyed multiple marmoset brain regions, including motor cortex, somatosensory cortex, visual cortex, striatum, hippocampus, etc.

### STARmap procedure

The STARmap PLUS procedure was conducted following established protocols. Glass bottom 6-well plates were treated with methacryloxypropyltrimethoxysilane (Bind-Silane) and subsequently treated with poly-D-lysine solution. #2 Micro cover glasses (18 mm) were pretreated with Gel Slikck according to the manufacturer’s instructions. 20 μm coronal brain slices were mounted in the pretreated glass-bottom 6-well plate. The brain slices were fixed with 4% PFA in PBS at r.t. for 15 min, permeabilized with pre-chilled methanol at -80°C for 2hr, and rehydrated with PBSTR/Glycine/YtRNA at room temperature for 10 min before hybridization. SNAIL probes were dissolved at the concentration of 50 nM per probe in water, and the final concentration per probe for hybridization was 5 nM. The brain slices were incubated in 600 μL hybridization buffer (consisting of 2X SSC [Sigma-Aldrich, S6639], 10% formamide [Calbiochem, 655206], 1% Triton X-100, 20 mM RVC [Ribonucleoside vanadyl complex, New England Biolabs, S1402S], 0.1 mg/ml yeast tRNA, 0.5% SUPERaseIn, and SNAIL probes) at 40°C for 36-40 hours with gentle shaking. Subsequently, the samples were washed at 37°C for 20 min with 1200 μL PBSTR (PBS, 0.1% Tween-20, 0.1 U/μL SUPERaseIn) twice and then washed once at 37°C for 20 min with 1200 μL High Salt buffer (PBSTR, 4XSSC). After a brief rinse with PBSTR at r.t, the samples were incubated for two hours with 600 μL T4 DNA ligase mixture (containing 0.1 U/μL T4 DNA ligase [ThermoFisher, EL0011], 1X T4 ligase buffer, 0.2 mg/mL BSA, 0.2 U/μL of SUPERase-In) at room temperature with gentle shaking. This was followed by two washes with 1200 μL PBSTR. Next, the samples were incubated with 600 μL of rolling-circle amplification (RCA) mixture (including 0.2 U/μL Phi29 DNA polymerase [Thermo Scientific, EP0094], 1X Phi29 reaction buffer, 250 μM dNTP mixture, 0.2 mg/mL BSA, 0.2U/μL of SUPERase-In and 20 μM 5-(3-aminoallyl)-dUTP [Invitrogen, AM8439]) at 4°C for 30 minutes and at 30°C for two hours. The samples were then washed twice with 1200 μL PBST (PBS, 0.1% Tween-20) and treated with 800 μL of 20 mM Acrylic acid NHS ester (Sigma-Aldrich, 730300-1G) in 100 mM NaHCO_3_ (pH 8) for one hour at room temperature. After a brief wash with 1200 μL PBST, the samples were incubated with 800 μL monomer buffer (containing 4% acrylamide, 0.2% bis-acrylamide, 2X SSC) for 30 min at room temperature. The buffer was removed, and 50 μL of polymerization mixture (0.2% ammonium persulfate, 0.2% tetramethylethylenediamine in monomer buffer) was added to the center of the sample. The samples were immediately covered by Gel Slick-coated coverslip and incubated for one hour at room temperature under nitrogen gas atmosphere. Following polymerization, the samples were washed twice with 1200 μL PBST for 5 min each. The tissue-gel hybrids were then digested with Proteinase K (Invitrogen, 25530049, 0.2 mg/ml in 50 mM Tris-HCl 8.0, 100 mM NaCl, 1% SDS) at room temperature overnight and then washed with 1200 μL of 1 mM AEBSF in PBST (Sigma-Aldrich, 101500) once at room temperature for 5 min, followed by two additional washes with PBST. The samples were stored in PBST at 4°C until imaging and sequencing. Before SEDAL sequencing, the samples were washed twice with the stripping buffer (60% formamide and 0.1% Triton X-100 in water) and treated with the dephosphorylation mixture (0.25 U/μL Antarctic Phosphatase [NEB, M0289L], 1X reaction buffer, 0.2 mg/mL BSA) at 37°C for one hour. Each cycle of SEDAL sequencing began with two washes using the stripping buffer (10 min each) and three washes with PBST (5 min each). For the six round of 461-gene sequence, the sample was then incubated with the “sequencing by ligation” mixture (0.2 U/μL T4 DNA ligase [ThermoFisher, EL0011], 1X T4 ligase buffer, 0.2 mg/mL BSA, 10 μM reading probe, and 300 nM of each of the 16 two-base encoding fluorescent probes) at room temperature for three hours. After three washes with wash and imaging buffer (10% Formamide, 2X SSC in water) and DAPI staining, the sample was imaged in the wash and imaging buffer.

### Image processing

Images were deconvoluted by Huygens Essential version 21.04 (Scientific Volume Imaging, The Netherlands, http://svi.nl), using the CMLE algorithm, with SNR:10 and 10 iterations. Image registration, spot calling, and barcode filtering were performed by following previously published reports^49^ with minor adjustments.

### Cell segmentation

A pretrained machine learning model from the StarDist package^50^ was used to automatically identify nuclei from the 2D maximum projection of the DAPI staining image. The segmented image was then used to extract cell locations and serve as markers for cell body segmentation. To represent cell bodies, an overlay of stitched DAPI staining and merged amplicon images from the first sequencing round was created. A gaussian filter with σ equal to 3 was applied to this composite image before binarizing it using Otsu thresholding strategy. To better incorporate amplicons around the peripheral region of cell bodies, a binary dilation with a disk structure element (r = 5) was applied on the mask. Finally, a marker-based watershed transform was performed on the binary mask representing cell bodies for segmentation purposes. Points overlapping each segmented cell region in 2D were assigned to that specific cell in order to compute a per-cell gene expression matrix.

### Data preprocessing

Normalization was performed using conventional single-cell methods in Scanpy: *sc*.*pp*.*normalize_total, sc*.*pp*.*log1p*, and *sc*.*pp*.*scale*, in that order, were applied to each dataset with default parameters.

### Smoothing

Data were spatially smoothed by performing the following: for each cell, spatial nearest neighbors were identified, the average expression vector within that neighborhood (including the given cell) was calculated, and the given cell’s expression vector was set to that average. Nearest neighbors were determined using scikit-learn’s *NearestNeighbors* model.

The subsampling approach was performed using numpy’s *random*.*choice* function with the random seed set to zero by default. We found that a ratio of 1:3 samples:neighbors tends to work well for most datasets. When applied to datasets of different resolution, the number of neighbors for each dataset was independently adjusted to achieve roughly the same length scale for each dataset. For our cross-technology demonstration, pairs of numbers of neighbors, *k*, and numbers of samples, *s*, were (30,10), (400,133), and (10,3) for STARmap, Slide-seqV2, and Visium datasets, respectively. Smoothing was then performed by setting the features of each cell to the average of their respective subsampled neighborhoods.

The concatenation approach was performed by identifying the neighbors of each cell in order of proximity, including the given cells themselves, and concatenating them in that order. The resulting feature vectors of each cell were of size *d*x*k*, where *d* is the number of genes measured and *k* is the number of neighbors identified for each cell.

### Clustering

PCA was performed using scikit-learn’s *PCA* model. Leiden and UMAP were applied using Scanpy’s *sc*.*tl*.*leiden* and *sc*.*tl*.*umap* functions. Subclustering was performed hierarchically by performing low resolution clustering to identify “level1’’ clusters (non-neuronal, excitatory neuron, inhibitory neuron) followed by separate clustering of each level1 cluster to identify level2 clusters. K-means was performed by first smoothing features as described above, without subsampling. Then PCA was performed on the smoothed features. Scikit-learn’s *KMeans* model was applied to the resulting PC space to identify region clusters. Molecular cell type and region annotation for resulting clusters were performed by referencing marker genes for each cluster as well as comparing to the Allen Brain Atlas^51^ for mouse and Riken’s Marmoset Gene Atlas^20,21^ for marmoset. The NMF approach was performed by subtracting the minimum expression value to make nonnegative, smoothing as described above without subsampling, and fitting scikit-learn’s *NMF* model to the non-negative, smoothed data.

### Integration

Molecular cell type integration was performed by applying PCA to each dataset independently followed by applying Harmony to the stacked PC representations of the datasets. Molecular region integration was performed by subsampling, smoothing, and applying PCA to each dataset independently followed by applying Harmony to the stacked PC representations. The resulting Harmony-aligned PC representation was then used as input to downstream analyses, such as clustering, DEG analysis, and trajectory inference.

### DEG analysis

DEG analysis was performed by applying Scanpy’s *sc*.*tl*.*rank_genes_groups* function with default parameters. These default parameters entail performing the following for each gene. Cells are first separated into “self vs. other” groups based on the grouping parameter. For instance, say we focus on a given molecular region to identify its unique marker genes. Then the grouping would be performed by region, and a t-test would compare the expression histogram between region A and all others, region B and all others, and so on. Genes with large z-scores resulting from this test associated with a given region would be considered markers for that region. The same approach was used to generate species-specific markers for each region. In that case, cells from a given region (including both species) were isolated and passed into *sc*.*tl*.*rank_genes_groups* with the grouping parameter set to species. Then t-tests would be used to compare mouse region A to marmoset region A, mouse region B to marmoset region B, and so on in order to find species-specific markers for each region.

### Trajectory inference

Cells belonging to the cortical layer regions of the spatially integrated marmoset and mouse data were first isolated from the full dataset. Diffusion maps was then applied to the subsampled, smoothed, Harmony-aligned PC representation of the cortical data using Scanpy’s sc.tl.diffmap function with default parameters.

### Molecular depth comparisons

The molecular depth distribution of a given gene was compared across species as follows. First, trajectory inference was performed as described above to calculate a molecular depth value for each cell. Then, 50 evenly spaced bins were created along the depth range. Each cell was then assigned to the bin corresponding to its molecular depth, and its non-negative expression value for the given gene was added to the bin. Expression values for each cell were made non-negative by simply subtracting the minimum value across all cells. The expression of the given gene was then divided by the number of cells within each bin, respectively, to give an average approximation to the intensity of the gene’s expression at each molecular depth. The top and bottom 5 bins were omitted due to noise, as few numbers of cells had very high or low depth values. This was performed for each species separately, yielding a molecular depth curve of the given gene for each species. The curves were then normalized by dividing each curve by its sum. Curves for the given gene were then compared across species by calculating the absolute value of the difference at each depth bin and summing to form a measure of total difference between species.

The molecular depth distribution of a given cell type was compared in a simpler manner. Because each cell had a depth value, cells of a given type and species were isolated and histograms for each species were calculated and compared using a two-sided Mann-Whitney U-test and *P<0.01 after Bonferroni correction for multiple comparisons.

**Supplementary Figure 1.**
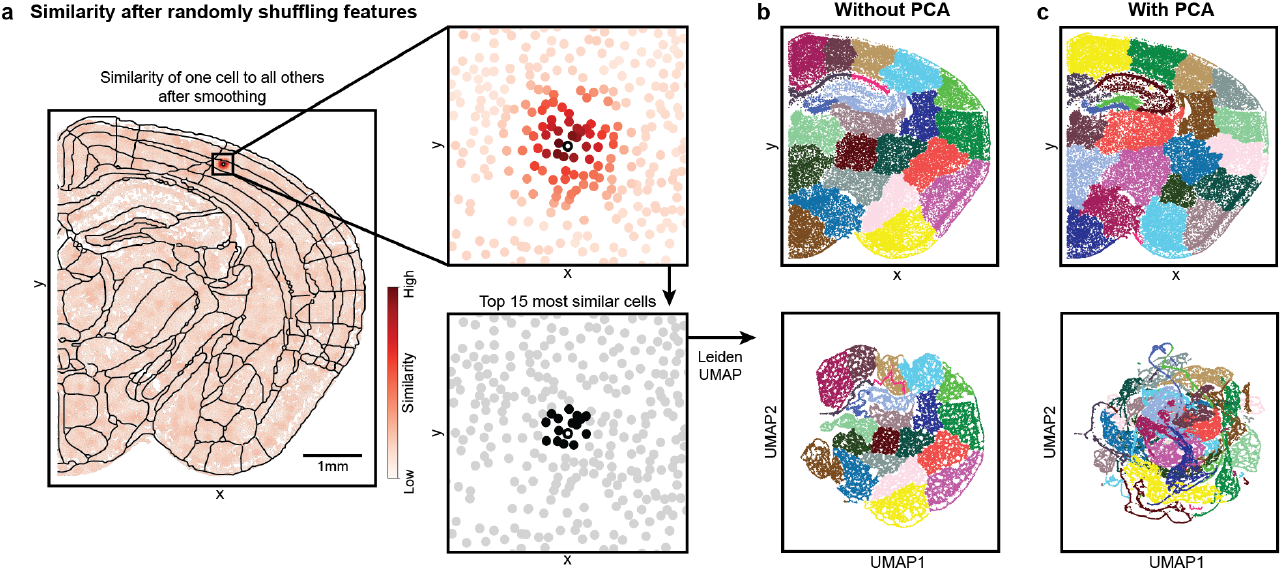
Spatial smoothing of randomly shuffled features yields spatial reconstruction. **a)** Results of comparing the smoothed representation of one cell (represented as a dot with white fill and black outline) to all others in the tissue. Similarity is defined as the dot product between normalized expression feature vectors. An anatomical wireframe from the Allen Brain Atlas is overlaid for comparison with conventional mouse brain anatomy^16^. Equivalent to Fig. 1d. **b)** Clustering results without using PCA, displayed in physical (above) and latent (below) spaces. Equivalent to Fig. 1e. **c)** Clustering results using PCA, displayed in physical (above) and latent (below) spaces.

**Supplementary Figure 2.**
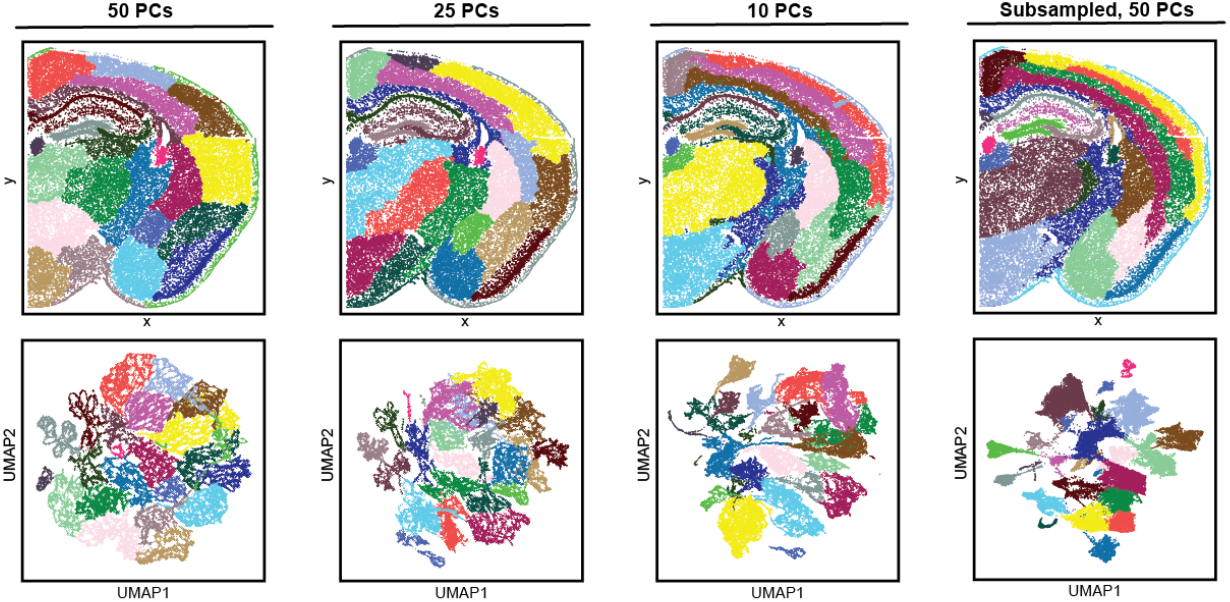
The effect of reducing the number of PCs used. Results from titrating the number of PCs used for Leiden and UMAP in the mouse brain STARmap PLUS sample. On the right, subsampling with 50 PCs is shown for comparison and is the same as Fig. 1h, Fig. 2c, and Fig. 3d. Leiden resolutions were set such that clustering yielded 23 clusters.

**Supplementary Figure 3.**
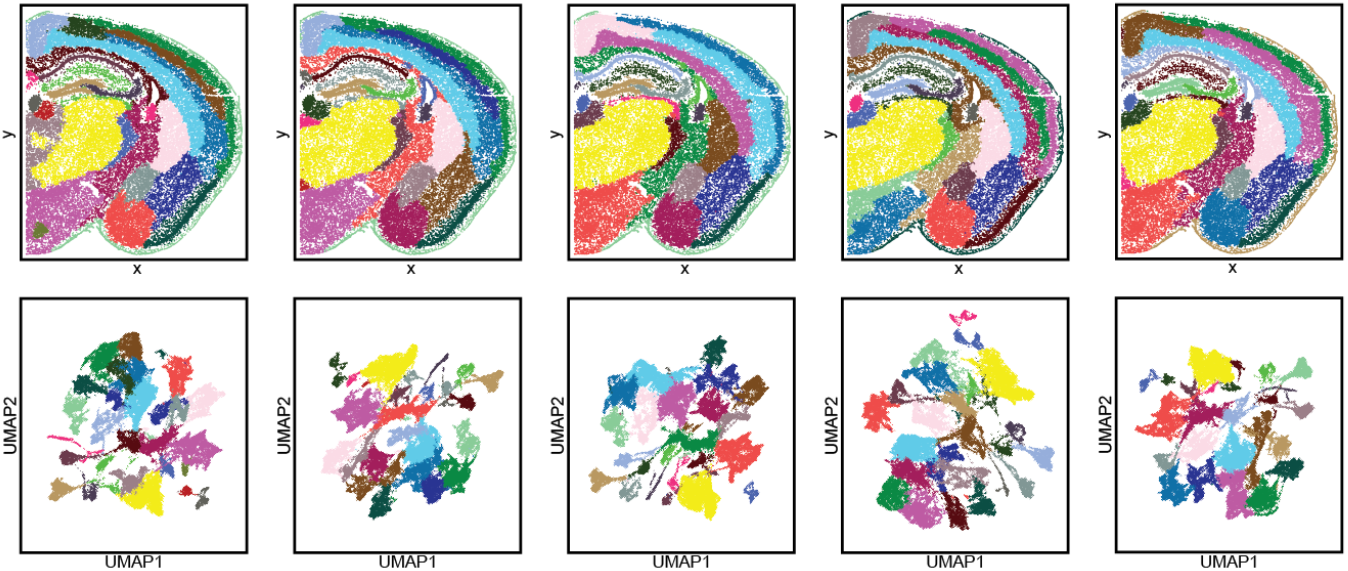
Consistency of SPIN across runs. Five runs of SPIN with different random states, each clustered using a Leiden resolution of 0.5. Each column represents the results of a separate run.

**Supplementary Figure 4.**
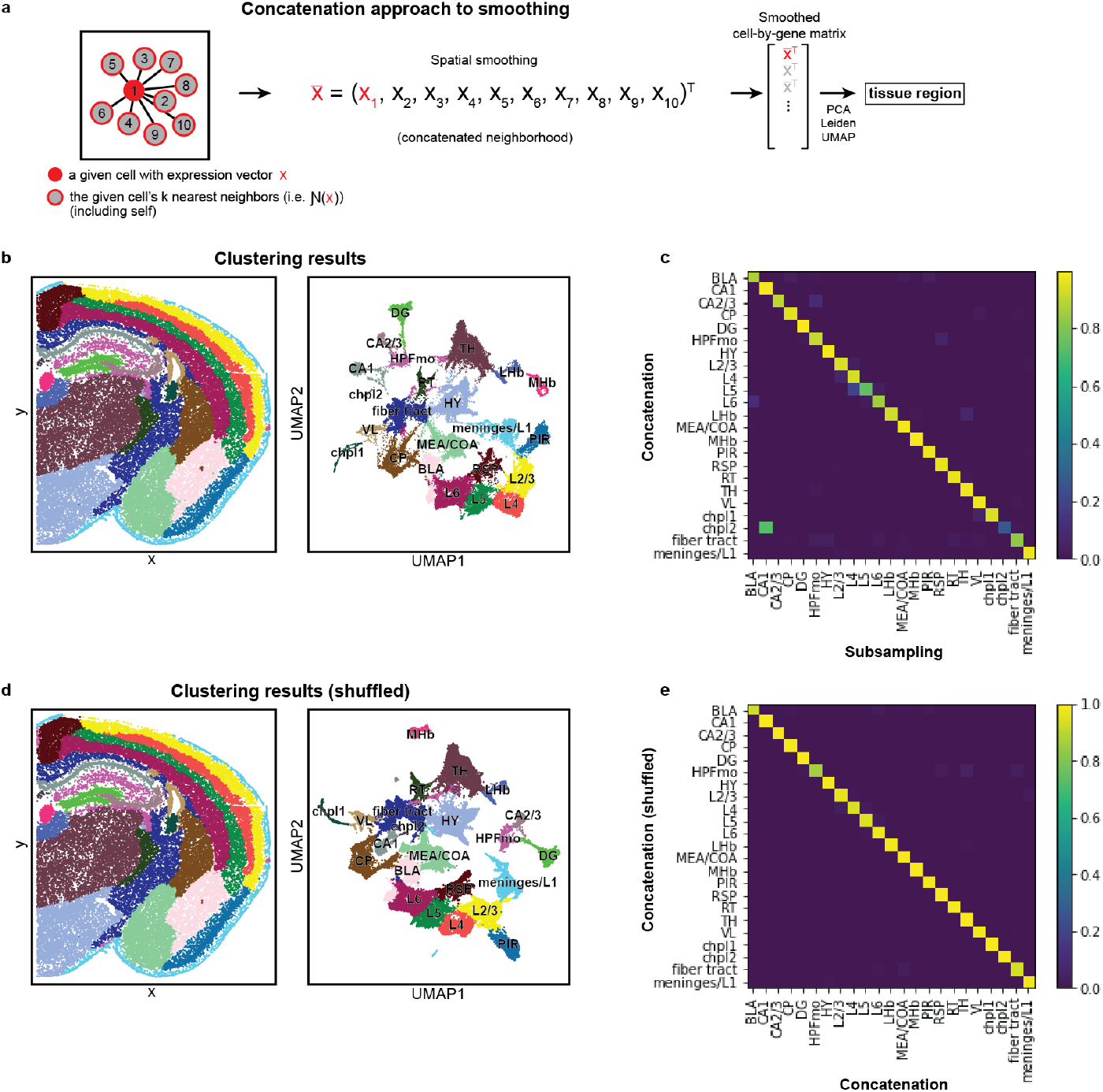
A concatenation approach to smoothing. **a)** Schematic of the concatenation approach. **b)** Results from clustering on the concatenated neighborhood representation. **c)** Confusion matrix representing comparison of cluster assignment between subsampling and concatenation approaches. **d)** Results from clustering on the concatenated neighborhood representation after shuffling the order of cells in each neighborhood. **e)** Confusion matrix representing comparison of cluster assignment between shuffled and plain concatenation approaches.

**Supplementary Figure 5.**
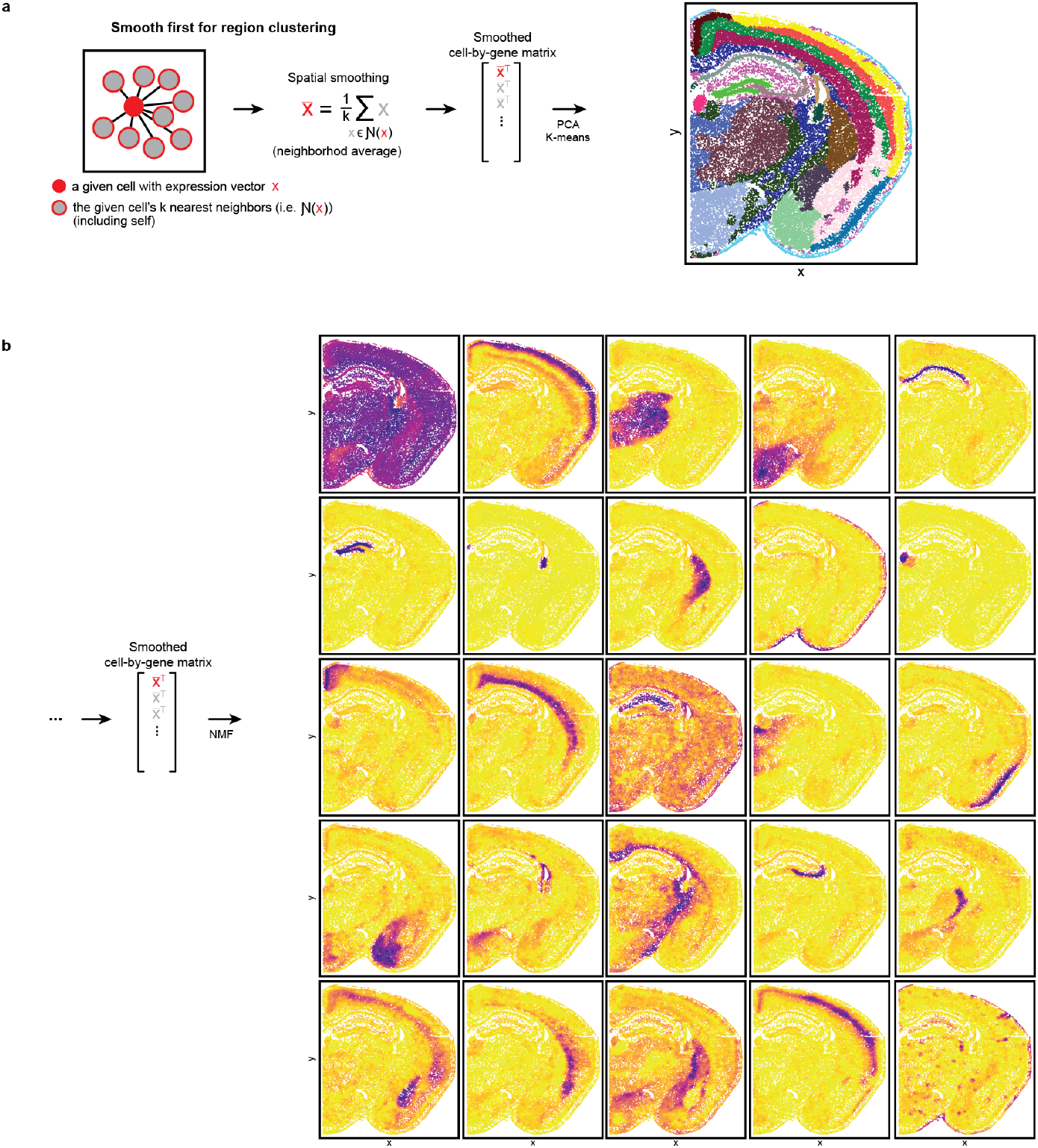
K-means and NMF approaches to smoothing avoid spatial reconstruction. **a**) K-means applied to spatially smoothed data with k=23 clusters. **b)** NMF applied to spatially smoothed data. Each of the resulting 25 factors is shown.

**Supplementary Figure 6.**
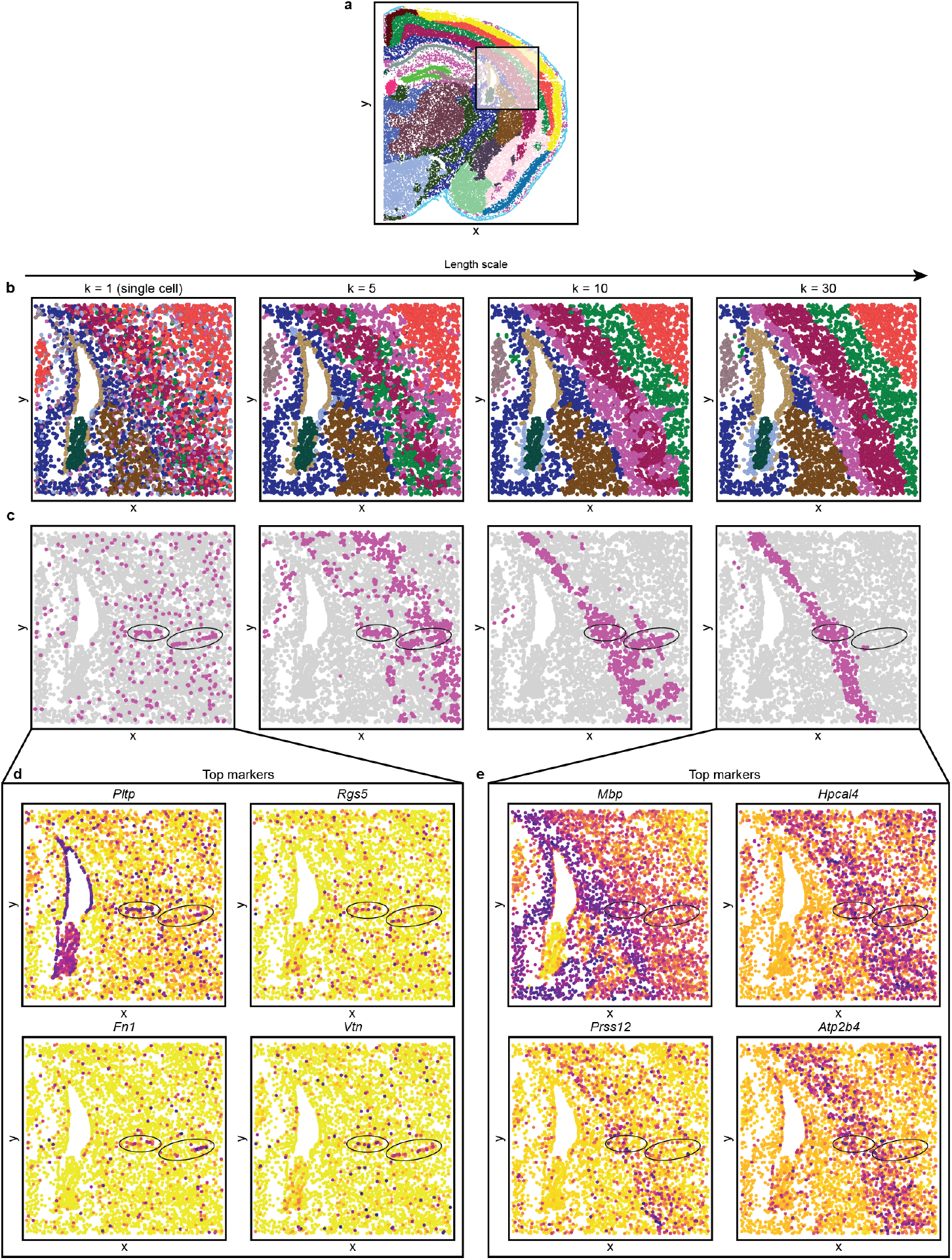
Titrating the neighborhood size, k, used for spatial smoothing. **a)** The region of interest used in b-e) **b)** K-means clustering results after spatial smoothing over various numbers of neighbors, *k*. K-means was used in order to avoid issues with subsampling on small length scales, i.e. for small *k*. **c)** Visualization of a cluster that spatially resembles vasculature after smoothing at lower length scales and L6b/fiber tract at higher length scales. Ovals outline vascular shapes in the *k*=1 clusters, which decrease in prominence at higher length scales. **d)** The top 4 marker genes of the vascular cluster at *k*=1. **e)** The top 4 marker genes of the L6b/fiber tract cluster at *k*=30.

**Supplementary Figure 7.**
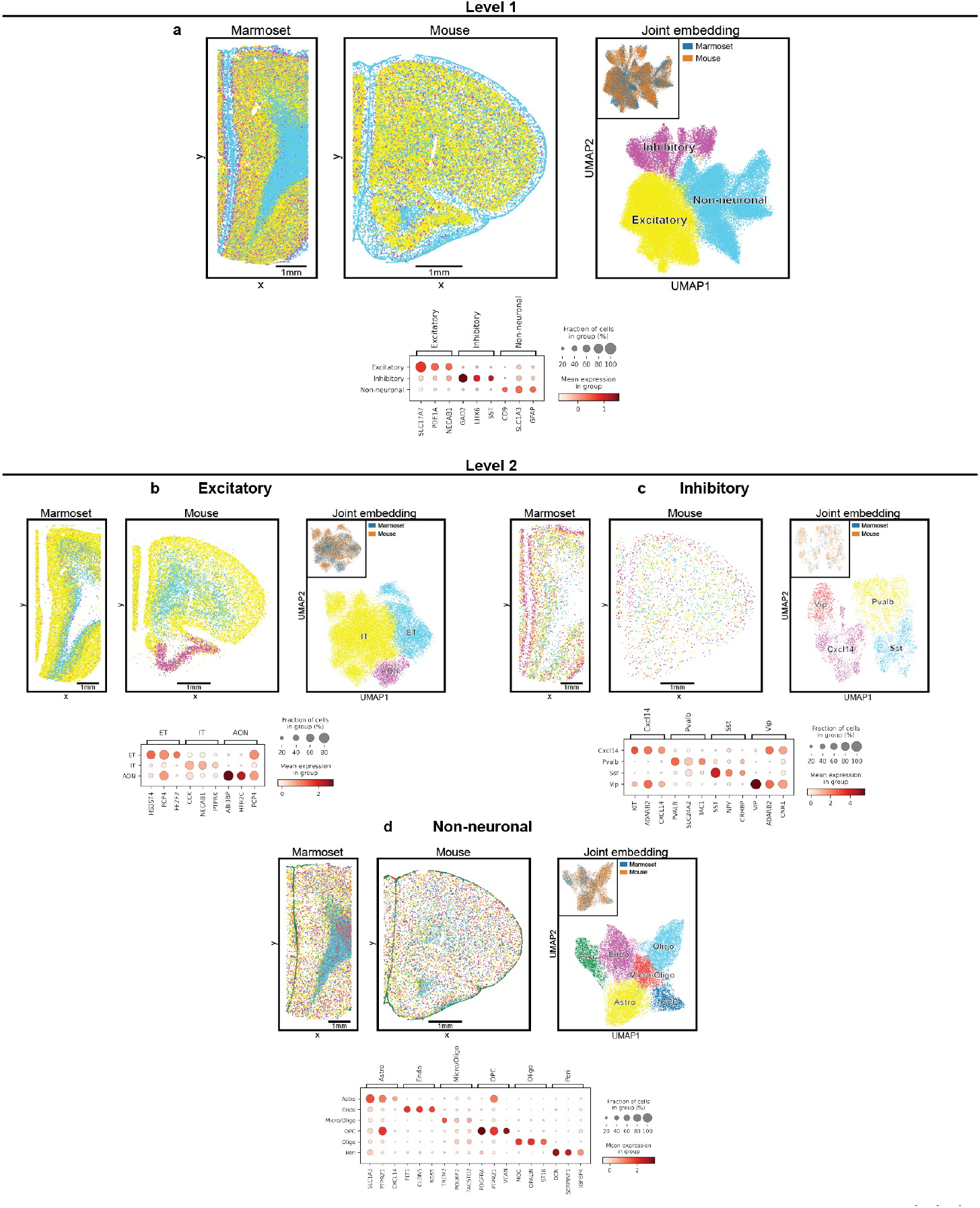
Subclustering results for molecular cell typing across species. **a)** Level 1 clustering results identify major brain cell type classes. Cell types plotted *in-situ* (above, left) and in UMAP space (above, right). Cells in UMAP space colored by species in upper inset (above, right). Top 3 marker genes for each cell type cluster (below) **b)** Results from subclustering the Excitatory neuron cell type cluster. Layout same as a).**c)** Same as b) but for the Inhibitory neuron cluster. **d)** Same as b,c) but for the Non-neuronal cluster.

**Supplementary Figure 8.**
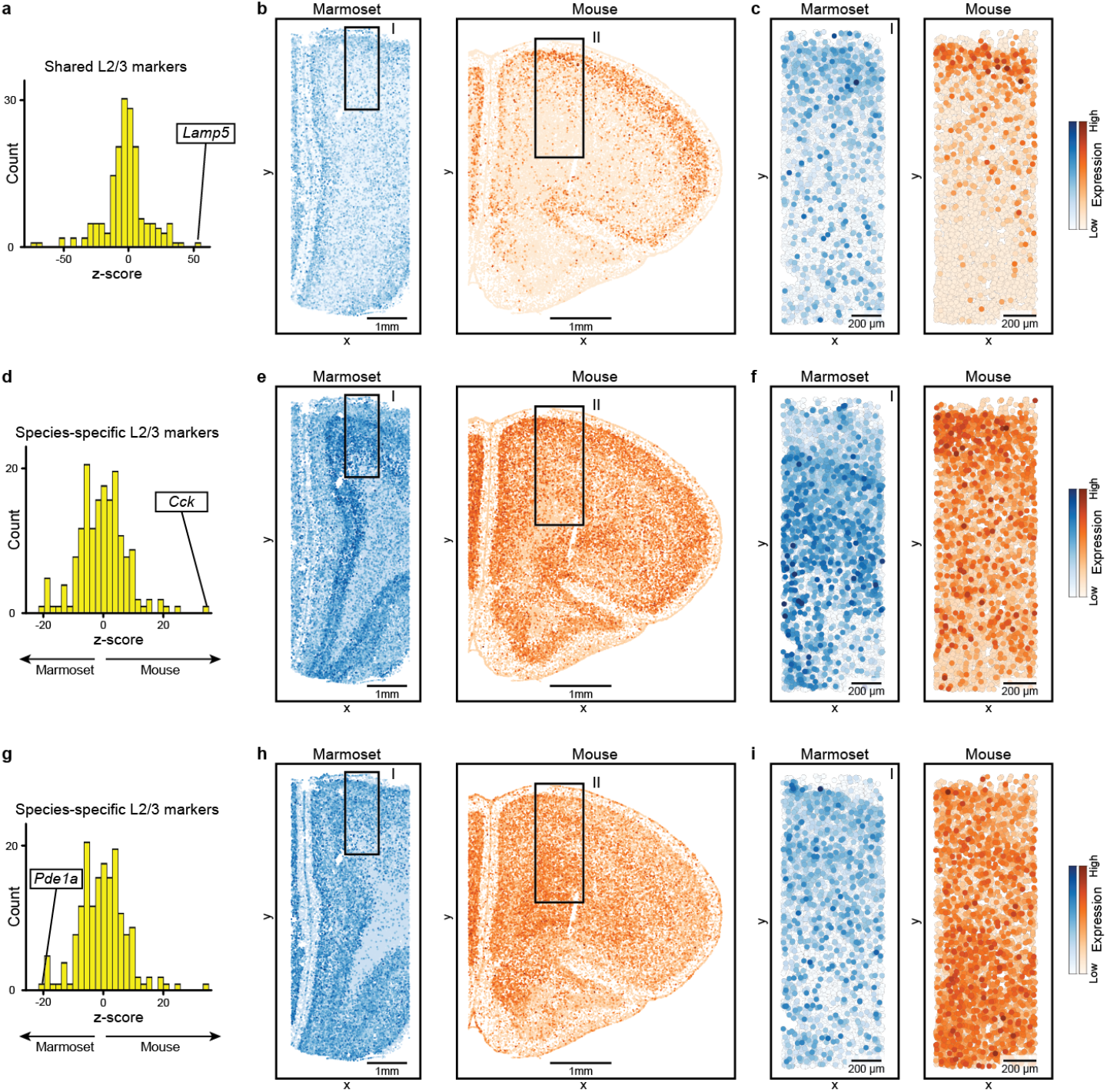
Visualization of shared and species-specific L2/3 marker genes. **a)** Histogram of shared L2/3 marker scores for each gene. Same as Fig. 4g (left). **b)** *Lamp5* expression plotted using the full tissue samples. **c)** *Lamp5* expression plotted on tissue zoom-ins. Same as Fig. 4g (right). **d)** Histogram of species-specific L2/3 marker scores for each gene. Same as Fig. 4h (left). **e)** *Cck* expression plotted using the full tissue samples. **f)** *Cck* expression plotted on tissue zoom-ins. Same as Fig. 4h (right). **g-i)** Analogous to d-f) but showing *Pde1a*. Colormaps were normalized for each marker gene independently.

